# Preserved Neural Dynamics across Arm and Brain-controlled Movements

**DOI:** 10.1101/2025.11.27.691057

**Authors:** Chenyang Li, Xinxiu Xu, Tianwei Wang, Yun Chen, Cong Zheng, Yiheng Zhang, Qifan Wang, He Cui

**Affiliations:** Beijing Institute for Brain Research, Chinese Academy of Medical Sciences & Peking Union Medical College, Beijing, 102206, China; Chinese Institute for Brain Research, Beijing, 102206, China; Center for Excellence in Brain Science and Intelligence Technology, Institute of Neuroscience, Chinese Academy of Sciences, Shanghai 200031, China

## Abstract

The neural activity for motor control is complex and dynamic; it has been found to dramatically transit from planning to executing movements. As brain-machine interfaces (BMIs) can directly connect the brain and the external world by yielding comparable motor outcomes with artificial apparatus, a central question is whether the BMI-controlled movements share neural dynamics or underlying mechanism with natural movements. To enable a systematic comparison, we developed a feedforward BMI framework with distinct planning and executing epochs that enables ballistic cursor control to intercept moving targets. This BMI allowed monkeys to voluntarily initiate neural states which controlled the direction and timing to launch a ballistic movement, like skeet shooting. Based on this, we found similar neural representations and computational structures across arm- and brain-controlled conditions. Notably, in addition to resembling the rotational structure in natural reaching, the neural population dynamics during open-loop BMI also shared preserved manifolds with those during reaching arm movements. These findings suggest a fundamental principle, and reveal a set of basic computational motifs for the neural control of movement in an abstract hierarchy in the absence of constraints from actuators. This study thus has the potential to reshape the consideration of how BMIs assist paralyzed patients in interacting with dynamic environments, and to promote next-generation BMI systems.

## Introduction

A brain-machine interface (BMI) enables an alternative and direct way for the brain to interact with the external world. By decoding neural signals in real-time, BMIs have succeeded in replacing lost motor functions (Bensmaia & Miller, 2014; Gilja et al., 2012; Lebedev & Nicolelis, 2017; Pandarinath & Bensmaia, 2022; Velliste et al., 2008). In recent years, a growing body of work has shown that BMI control recruits many sensorimotor brain-regions engaged during able-bodied movement (Golub et al., 2015; Stavisky et al., 2017). These parallels support the notion that high-performance BMIs may rely on sensorimotor processes similar to those underlying natural motor control (Stavisky et al., 2017), thereby motivating biomimetic BMI design. Nevertheless, direct evidence that the neural mechanisms underlying sensorimotor processes are preserved between natural movement and BMI control is still lacking.

Addressing this question requires a theoretical lens capable of capturing the essence of motor control. For cortical movement control, recent decades have seen a shift from seeking explicit movement-parametric representations to adopting a population-level, dynamical-systems framework (Vyas et al., 2020). This has explained once-puzzling phenomena, such as the dramatic shift in neural dynamics of the motor cortex when motor preparation switches to execution (Churchland & Shenoy, 2024; Kaufman et al., 2014). In this view, the motor cortex acts as a pattern generator whose neural dynamics are initialized by preparatory activity and evolve during motor execution (Shenoy et al., 2013). Despite its explanatory power, however, these fundamental principles remain underutilized. While models based on this perspective have improved decoding by focusing on state evolution rather than parameter estimation (Kao et al., 2015; Pandarinath et al., 2018), the dominant paradigm for current BMIs, iterative feedback control, is distinct from the autonomous evolution envisioned by this dynamical framework, and may thus inherently impose a performance-ceiling. Therefore, a critical opportunity lies in implementing a biomimetic BMI that is compatible with this framework to directly test whether neural dynamics are preserved during brain control.

In the present study, we designed a feedforward BMI that enables monkeys to control a computer cursor to move with ballistic trajectories, allowing it to intercept rapidly moving targets. Such a BMI framework features well-defined preparation and execution epochs, via an epoch-switch detector that allows self-paced movement initiation. Strikingly, neural responses during this biomimetic brain-control exhibited the prominent dynamical characteristics observed during naturalistic arm reaching, and they shared a low-dimensional neural manifold with arm reaching, suggesting that similar neural dynamics underlie both brain-controlled and arm movements. Our study thus provides direct evidence that neural dynamics are preserved across BMI and natural movement, offering not only novel insights for fundamental sensorimotor control but also a principled path toward next-generation BMIs.

### Feedforward BMI for intercepting moving target

Given similar task demands, it is reasonable to expect that a monkey would use similar low-dimensional neural manifold to finish the interception for both BMI and natural movements, constrained by basic neural circuits and behavioral strategy (Perich et al., 2025). We hypothesize that, with appropriate biomimetic design, the latent states for both BMI and natural reaching would follow similar traces on the shared preparatory and execution subspaces (**Fig. 1a**), reflecting the characteristic initial-state-dependent evolution of motor cortical dynamics (Shenoy et al., 2013).

**Figure 1.**
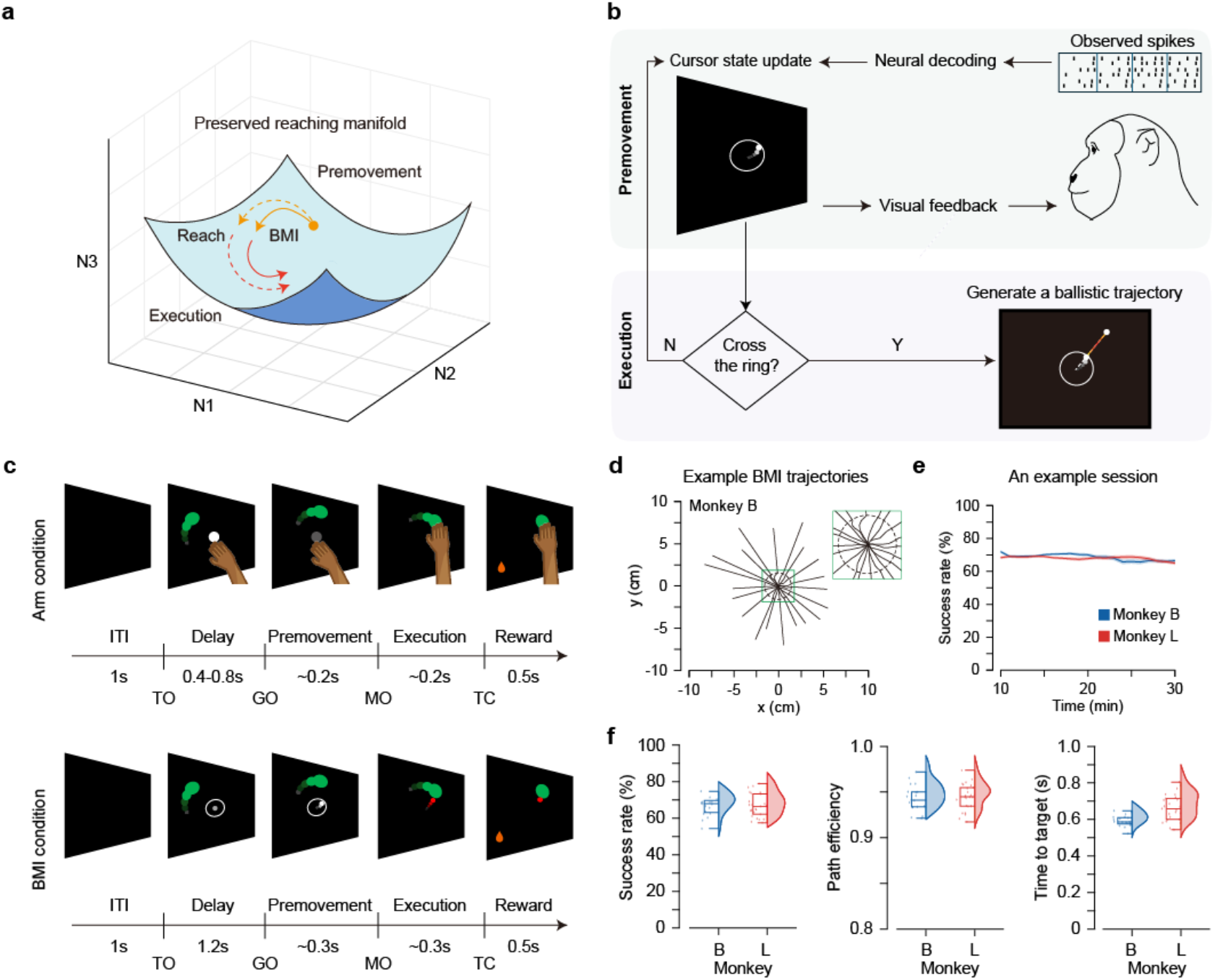
The feedforward BMI and its performance. (**a**) Hypothesis for preserved neural dynamics across brain-controlled and arm movement. We supposed that the reaching-like dynamics would emerge with the explicitation of feedforward BMI control due to the intrinsic properties of the motor cortex. (**b**) The schematic diagram of the feedforward BMI framework, featured by distinct premovement and execution epochs. In the former epoch, the cursor is brain-controlled to exceed a threshold in a closed-loop manner. Once the cursor crosses the threshold, the movement course switches to the next epoch and the cursor launches ballistically in an open-loop manner. (**c**) The interception task including arm and brain-controlled conditions. TO: target on, GO: GO-cue, MO: movement onset, and TC: touch target. (**d**) Example BMI trajectories of monkey B. The dotted ring indicates the ring to be crossed to trigger movement. The inset zooms in the situation inside the ring. (**e**) Success rates over the course of an example session. Data are excluded from the initial 10-minute decoder calibration period. The trace shows performance calculated with a sliding window (width: 75 trials; stride: 1 trial). (**f**) Averaged success rate, path efficiency, and time to target of BMI sessions. The success rate for monkey B was 66.8±2.6% across 19 sessions and for monkey L 66.9±2.6% across 25 sessions (mean ± 1.96 s.e.m.). The path efficiency was 0.94±0.01 for monkey B and 0.96±0.01 for monkey L. The time to target (interval from the cursor initiation to target acquisition) was 0.59 ± 0.02 s and 0.66 ± 0.03 s, comprising approximately 0.31 ± 0.01 s and 0.37 ± 0.03 s in the premovement epoch and 0.28 ± 0.01 s and 0.30 ± 0.01 s in the execution epoch for Monkeys B and L, respectively.

To test this hypothesis, we designed a feedforward BMI framework with the following key features (**Fig. 1b**): (1) Clearly delineated preparation and execution epochs. During the preparation epoch, a decoder enables subjects to initialize neural states explicitly with visual feedback. During the execution epoch, the initial state (i.e. the preparatorily set state) determines the launch direction of a ballistic movement with a bell-shaped speed profile in a feedforward manner. Additional feedback-based adjustments, implemented using the same decoder, can be incorporated to provide online corrections if necessary; (2) A detector or “gate” separates the two epochs and acts as an onset trigger of ballistic movement. Crucially, the gate’s activation is determined by the subject’s own intent, rendering the BMI asynchronous, as movement initiation is detected directly from neural intention.

As an instantiation of such biomimetic BMI framework, we recorded neural population activity from two microelectrode arrays implanted in the primary motor cortex (M1) and dorsal premotor cortex (PMd) of monkeys B and L (**Fig. S1**). Monkeys were firstly trained to manually perform a delayed center-out reach task on a vertical touchscreen displaying static or moving targets, which is referred as the manual task or arm condition (**Fig. 1c**). Previous studies have shown that intercepting moving targets requires precise spatiotemporal coordination, making it a suitable task for evaluating BMI performance (Li et al., 2018; Zhang et al., 2025). We used a target speed of ±120 °/s in recording sessions, which is appropriate for most monkeys. For the BMI condition (**Fig 1c**), we first calibrated a velocity decoder using threshold-crossing firing rates recorded while the monkey passively observed a computer-controlled cursor performing center-out movements. Then the monkey, with its arms restrained in the chair, was trained to control the cursor to touch a static target in a closed-loop decoder adaptation manner (Gilja et al., 2012; Orsborn et al., 2014) until computer assistance was removed and performance exceeded 80% success. Next, we introduced a white ‘indicator ring’ around the center. After a 1-sec delay period in which target was displayed, the monkeys were allow to brain-control the cursor from the center. Once the cursor crossed the ring, it was launched centrifugally along the radial direction of the crossed point, following a straight trajectory with a fixed bell-shaped speed profile, whose initial velocity matched the cursor speed at ring crossing (example trajectories in **Fig 1d**). We defined the period during which cursor remained inside the ring as the “premovement” epoch, corresponding to preparation period before GO in the manual task. If the cursor entered a 5-cm radius around the target, a reward was delivered. If the cursor overshot and missed the target, the monkey was allowed to control the cursor using visual feedback. If the target was not reached within the following 1 s, the trial was marked as a failure, and no reward was given.

After few weeks of training, both monkeys successfully learned to perform the BMI task with the feedforward strategy during comparable time course (**Fig. S2a, d**). Task performance remained stable throughout each session (**Fig. 1d**), with mean success rates near 70% (**Fig. 1e**). In most of the successful trials, the cursor traveled directly toward the target without requiring any feedback-based corrections. As in the manual task, movement trajectories were uniformly distributed across directions in the BMI task without directional bias (**Fig. S2b-c, e-f**). For subsequent analyses, trials were grouped into eight reach directions evenly spaced at 45° intervals.

### Across-epoch change in neural representation

To investigate whether the feedforward BMI task resembles the manual task, we first analyzed single-unit responses in the different datasets. As in the manual task, neurons during the BMI task also exhibited sequence-like firing patterns characteristic of natural reaching, albeit with reduced temporal sparsity (**Fig. S3**). We fit each unit’s activity with a cosine tuning model and quantified the percentage of units with a goodness-of-fit greater than 0.7 (tuned units’ percentage, TU%). During the inter-trial interval (ITI), TU% of was near zero for both tasks (**Fig. 2a**). After target onset, TU% began to rise in both tasks, though the increase in the BMI task was delayed and slower compared with the manual task, indicating a gradual engagement of neural population. During the pre-movement epoch, TU% reached a plateau that was comparable across tasks for monkey B but slightly lower in the BMI task for monkey L, probably reflecting sampling bias in recorded neurons. Notably, during the execution epoch when the cursor moved without direct brain control, TU% in the BMI task remained similar to that in the manual task (51.2 ± 3.1% for monkey B; 44.3 ± 2.5% for monkey L). To rule out the effect of visual feedback, we further occluded the cursor during its flight (**Fig. S4a**), so that it appeared to the monkey that the cursor jumped directly from the center to the periphery. Even if the cursor was invisible in-flight, TU% remained high in the execution epoch (**Fig. S4c**). Given that the cursor motion directly determined reward outcome and that monkeys were afforded an opportunity to correct potential target misses, the sustained TU% observed during the execution epoch suggests that monkeys were still actively engaged in the open-loop control, rather than passively observing the cursor’s ballistic motion.

**Figure 2.**
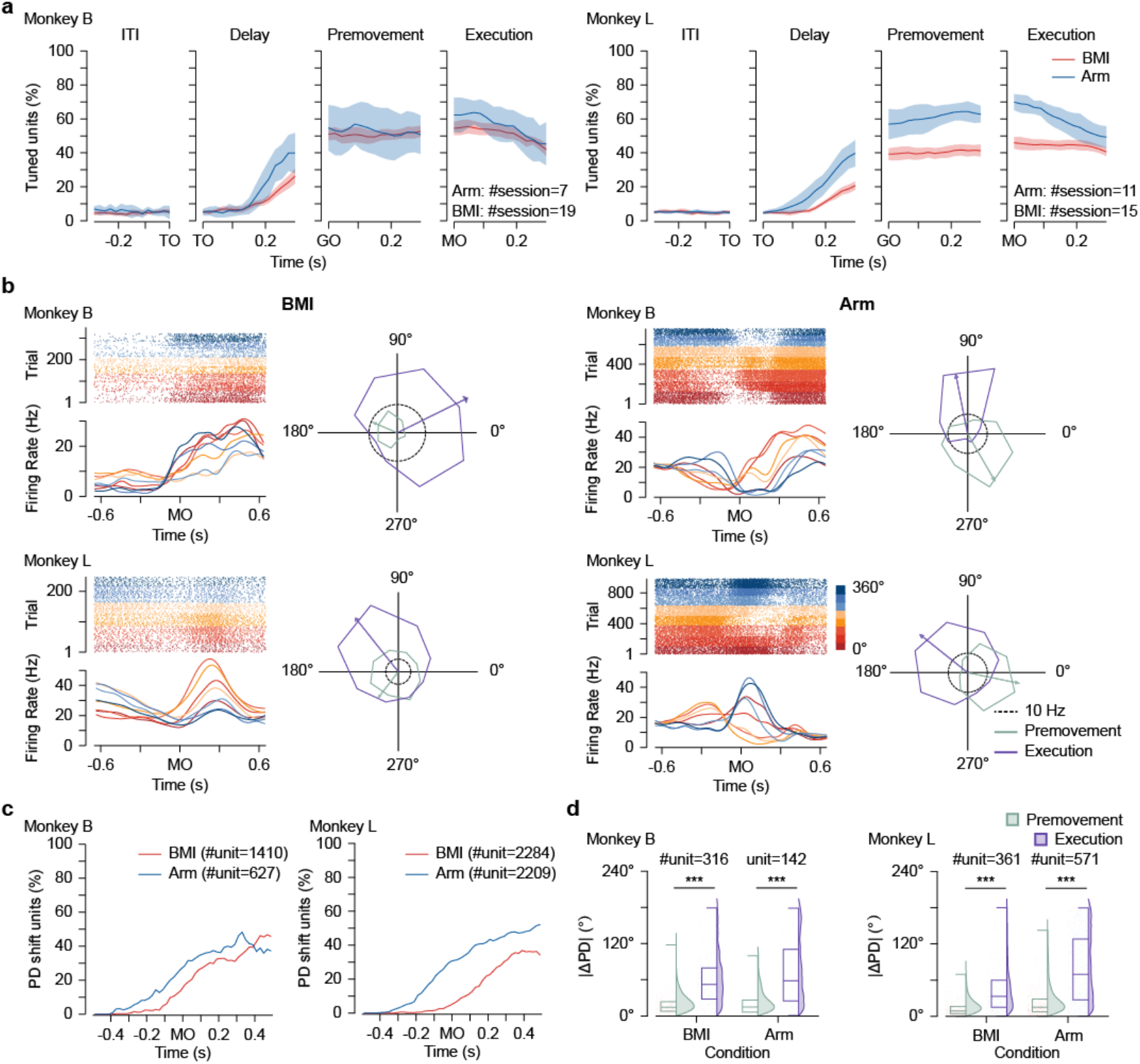
Single-neuron tunings to arm- and brain-control movements. (**a**) Tuned unit’s percentage (TU%) in different epochs when aligned to target on (TO), Go-cue on (GO), and movement onset (MO) in the BMI (red) and arm (blue) conditions; (**b**) Example neurons with PD shift in the BMI and arm conditions. For each subplot, on the left is the raster plot and peri-event time histogram (PETH), aligned to MO and colored by movement direction; on the right is a polar plot, where the angle denotes the movement direction, the radius denotes the averaged firing rate in the corresponding direction, and the arrow shows the preferred direction (PD) in the certain epoch. Premovement epoch: −200 ms to −180 ms from MO; Execution epoch: 180 ms to 200 ms from MO. (**c**) The proportion of ‘PD shift’ neurons over time relative to MO. Criteria for ‘PD’ neurons: goodness-of-fit of the cosine tuning model was greater than 0.7 both in 0.5 s before MO and the current time. Criteria for ‘PD shift’ neurons: the difference between current PD and the PD in 0.5 s before MO exceeded 90°. (**d**) Absolute differences of PD between premovement and delay epochs (green) and between execution and delay epochs (purple) in both conditions.

We further characterized the neural representation pattern by computing each neuron’s preferred direction (PDs, (Georgopoulos et al., 1982) and comparing dynamic changes of PD under two conditions. During natural reaching, motor cortex neuron could have two stable and distinct PDs during preparatory and execution periods (Suway et al., 2018). Here, we defined a neuron as exhibiting a “PD shift” if the angular difference between PDs of two epochs exceeded 90°. As expected, neurons in both tasks exhibited varying degrees of PD shift from premovement to execution epoch (**Fig. 2b**). Changes in modulation depth and baseline from the pre-movement to execution epochs were also comparable across tasks, further supporting a similar across-epoch representational transition (**Fig. S5**). The proportion of PD-shift neurons in the tuned units increased over time (**Fig. 2c**). In the manual task, this increase began ∼0.4 s before movement onset (MO), whereas in the BMI task the ramping began 0.1-0.2 s later. The overall fraction of PD-shift units was consistently lower in the BMI task than in the manual task (execution-epoch plateau: 36.1% in manual task and 32.7% in BMI task for monkey B; 43.4% and 22.7% for monkey L, from perimovement to execution epoch). As expected for a continuous preparatory process, the PD shift from the delay to premovement epoch in the manual task was smaller than the PD shift from delay to execution epoch (**Fig. 2d**). Strikingly, this relationship was preserved in the BMI task. This indicates that the delay epoch when cursor is static and pre-movement epoch when cursor was controlled form a continuous neural process, whereas the transition from pre-movement to execution epoch reflects a phase transition in neural dynamics.

### Neural population dynamics in arm and BMI tasks

Given the similar single-unit firing patterns observed across tasks, we next asked whether our feedforward BMI also reproduces the population-level dynamics characteristic of natural arm movements. In three-dimensional subspaces constructed from trial-averaged population data using principal component analysis (PCA), neural trajectories from the BMI and manual task exhibited two common features: first, the neural trajectories were ordered according to movement direction; second, although trajectories initially evolved outward from the origin, they reversed direction inward shortly after MO, transitioning from the premovement to the execution epoch (**Fig. 3a**). Examining individual principal components (PCs) revealed that this trajectory reversal at MO was primarily driven by PC3, which showed a value flip for all movement directions, whereas PC1 and PC2 remained relatively stable over time (**Fig. S6**). The shared neural geometry verified the PD-shifts observed at single-unit level and further suggested that our feedforward BMI preserves a key computational structure underlying natural arm movements.

**Figure 3.**
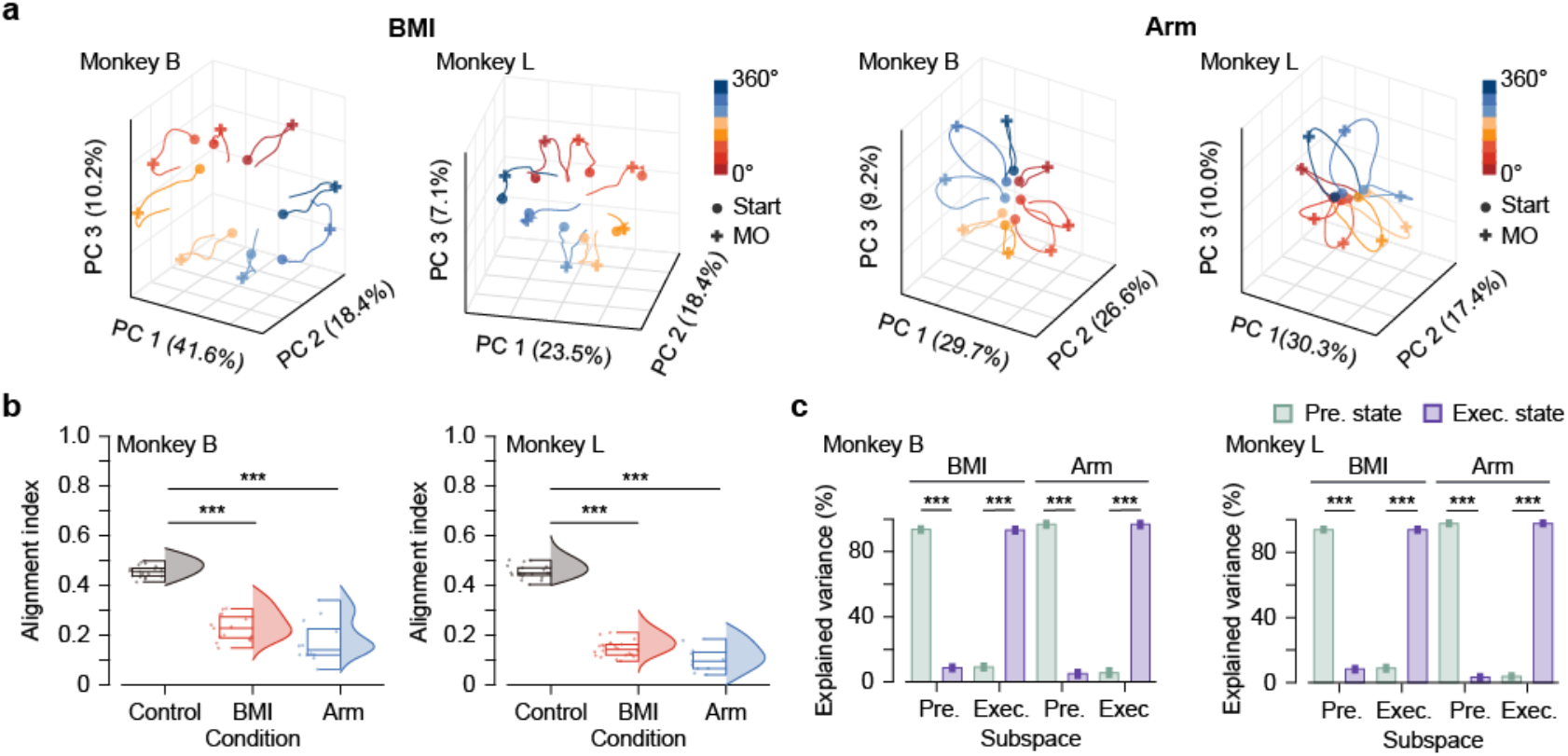
Neural state transition from the premovement to execution epochs revealed by dimensionality reduction. (**a**) Neural trajectories of the top three principal components (PCs) identified with principal component analysis (PCA) in the BMI and arm condition. Different colors represent distinct cursor’s ballistic directions. (**b**) Alignment index between premovement subspace and execution subspace under different conditions. Bars of ‘control’ correspond to the indices resulted from random dimensions. (**c**) Percentage of the premovement-epoch data variance and execution-epoch data variance explained by the top six execution-epoch PCs or premovement-epoch PCs in the BMI and arm conditions.

Orthogonality between preparatory and execution neural subspaces is another key feature observed during ballistic arm movements (Elsayed et al., 2016). To assess whether this property was also present in our BMI framework, we first quantified the extent of subspace overlap using the alignment index, defined as the fraction of execution-epoch variance captured by the premovement-epoch subspace spanned by the top ten PCs (Elsayed et al., 2016). Perfect orthogonality would yield an alignment index of zero. Consistent with previous findings, the alignment index for the manual task was near zero and significantly lower than the control level derived from simulated random data (**Fig. 3b**, 0.173 ± 0.053 for monkey B; 0.102 ± 0.033 for monkey L). As expected, the alignment indices of the BMI task were comparable to those of the manual task (0.228 ± 0.029 for monkey B; 0.145 ± 0.014 for monkey L). We further identified the ‘premovement subspace’ and ‘execution subspace’ and projected neural activity from each epoch into these subspaces, following prior methodology. In both the BMI and manual tasks, each subspace captured high variance within its corresponding epoch but minimal variance from the other epoch (**Fig. 3c**), indicating strong orthogonality between preparatory and execution dimensions. These results demonstrate that our feedforward BMI has well-defined preparatory and executing periods which recapitulate critical dynamical features observed during the ballistic arm movement.

### Preserved evolution rule and neural manifold

Neural population dynamics often evolve in a quasi-oscillatory manner across movement types, providing direct evidence for the computational mechanisms underlying cortical dynamics (Churchland et al., 2012). To examine whether this principle is also preserved in the feedforward BMI, we applied jPCA to neural activity pooled across sessions (−0.1 to 0.3 s around MO). Consistent with previous studies, neural dynamics in the manual task exhibited clear rotational structure (**Fig. 4a**). Remarkably, a similar rotational pattern was also present in the BMI task, indicating that the same underlying dynamical rule is preserved during biomimetic brain control.

**Figure 4.**
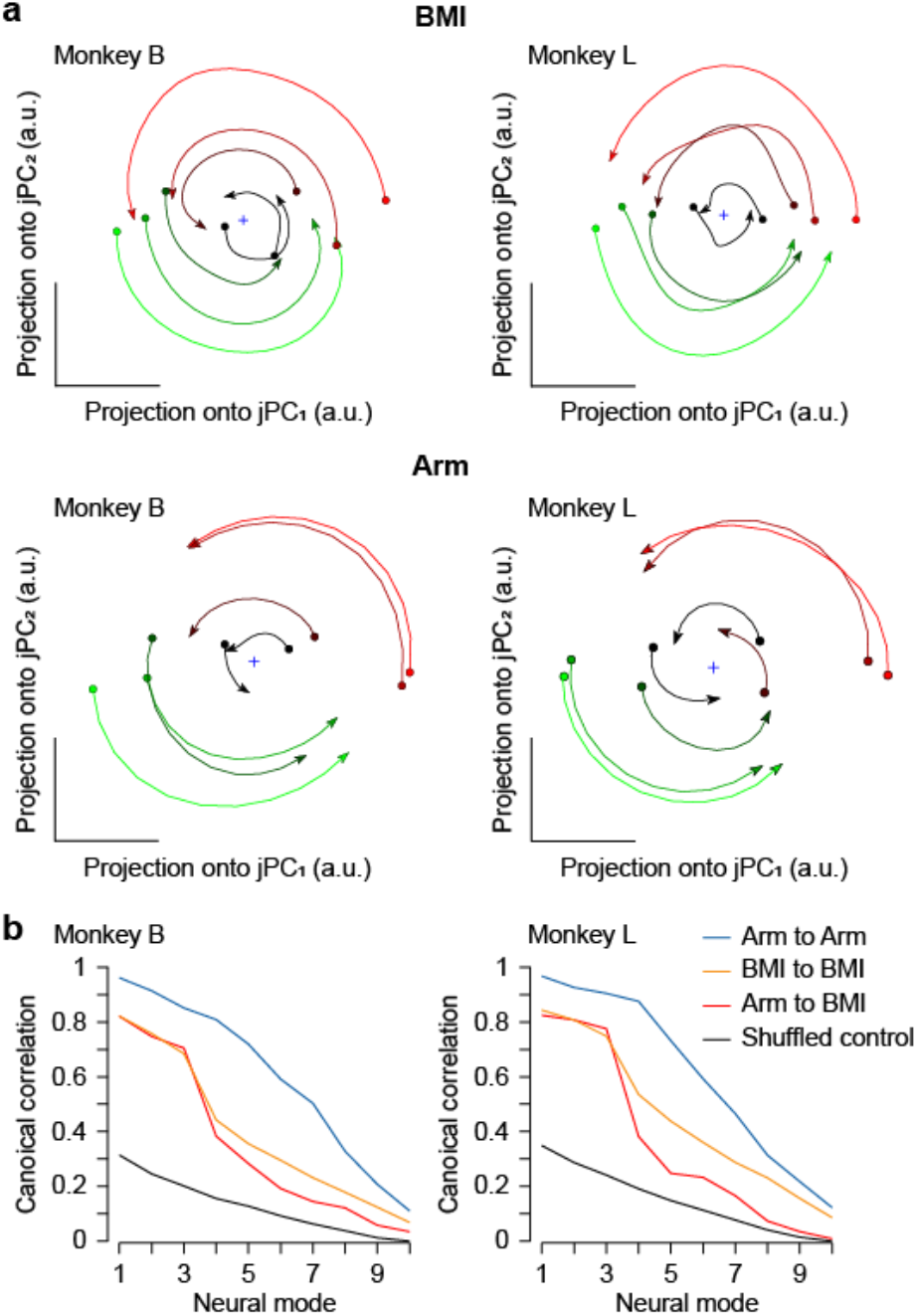
Neural latent dynamics during brain-controlled and arm movements. (**a**) Projection of neural activities to jPCA space in BMI and arm conditions. Each trajectory represents one of the eight movement directions. Trajectories are colored by their initial states. For monkey B, 486 neurons and 469 neurons were pooled in BMI and arm conditions, respectively; for monkey L, the numbers were 568 and 566. (**b**) Correlation changing in different neural mode. The correlation for aligned neural activity in PCA space were divided to four groups: within session of arm movement (blue), brain control (orange), across sessions (red), and random trials selected from all sessions (black). The time window was set to the 300 ms before the ballistic and movement onset.

Given the above findings, we finally asked whether the BMI and manual tasks shared a low-dimensional common neural manifold during the execution period. To identify potential shared neural modes, we performed canonical correlation analysis (CCA) on the top ten PCs of population data (−0.3 to 0.4 s around MO) from pairs of sessions, either from the BMI or manual task. Because neural modes were ordered by their corresponding PCs, canonical correlation naturally decreased with increasing mode index, reflecting diminishing explained variance (**Fig. 4b**). Neural modes from two manual-task sessions exhibited the highest correlations, whereas shuffled control constructed from simulated random data exhibited the lowest. Within-task correlations were lower for BMI sessions than for manual sessions, with the first three modes showed high correlation, and correlations for higher-order modes dropped sharply, suggesting that shared latent dynamics across BMI sessions were captured by fewer neural modes. Intriguingly, the correlation pattern between BMI and manual sessions resembled the within-BMI pattern, with the across-task correlations declining more steeply but remaining above the shuffled level. Together, these results provide direct evidence that the BMI and manual tasks shared a preserved low-dimensional neural manifold, despite their distinct effectors.

## Discussion

BMI might gain a decisive advantage by embracing well-established motor control principles, particularly frameworks that emphasize cortical preparatory dynamics in generating movements. The key insight is that by leveraging preparatory dynamics, control could be shifted forward in time, thereby potentially bypassing a fundamental source of system latency. Here, we proposed and implemented a biomimetic, feedforward BMI. It features a distinct preparatory stage, which optimizes the neural initial state and an autonomous execution stage where the cursor moves ballistically. This design successfully captured key dynamical features of natural arm movements, providing direct evidence for preserved neural dynamics across BMI and natural movements.

According to the internal model theory (Franklin & Wolpert, 2011), skilled movements are inversely preplanned via forward model of future states, rather than adjusted online during execution. We propose that the dynamical evolution from an initial state set by preparatory activity constitutes a plausible neural mechanism for such internal models and aligned our BMI design with this. Neural dynamics are known to preserve across effectors (arm or arm-control manipulandum), individuals, and species (Safaie et al., 2023; Stavisky et al., 2019). Our finding extends this conservation to the brain-controlled context. This consistency suggests a relatively abstract ‘motor protocol’ within the motor cortex. This protocol, independent of the specific actuator (arm or cursor), may implement a core component of the feedforward internal model. The required translation of this protocol into the actuator-specific commands thus reveals a critical axis in the hierarchy of motor control. Future work could “survey” this hierarchy by comparing the degree of preservation across biologically-defined cell types (Chen et al., 2023) or dynamical motifs (Driscoll et al., 2024). This hierarchy will likely deepen our understanding of sensorimotor control, and in turn, calls for the development of hierarchical decoding algorithms to orchestrate complex movements.

Improving BMI control of 3D robotic devices hinges on releasing the burden of calibration. While “manifold-alignment” algorithms address recording instability (Degenhart et al., 2020; Gallego et al., 2020; Karpowicz et al., 2025; Safaie et al., 2023), a parallel strategy seeks to build foundational models pre-trained on large-scale neural datasets (Ye et al., 2023; Zhang et al., 2024). However, both approaches confront a fundamental bottleneck: the scarcity of subject-specific neural data for training, which is especially prohibitive for patient populations. Our finding directly addresses this bottleneck, as it ensures that the brain operates in a familiar dynamic regime during BMI control. This justification enables a novel strategy: to leverage large-scale, able-bodied neurobehavioral datasets during natural movement to train a foundation model for BMI. This strategy can then enable rapid personalization and may serve as a starting point for building generative BMIs, which could produce fluent, context-appropriate motor programs from sparse neural ‘prompts’.

In summary, we introduced a biomimetic BMI and demonstrated preserved neural dynamics. This work provides not only a novel experimental framework for sensorimotor research but also a principled bridge to next-generation, generative BMIs.

## Methods

### Experimental preparation

Two male macaque monkeys (*macaca mulata*, B and L, weighting 5-9 kg) were trained to perform interception in both manual and brain-controlled interceptions. In each session, the animal was seated in a custom-made primate chair facing a vertical touchscreen (Elo Touchsystems, 19”; sampling at 60 Hz, spatial resolution <0.1 mm) 30 cm away. The animal was head-fixed in the primate chair via two headposts implanted into the skull along the midline suture. Hand positions were monitored by recording the location of an optically reflective marker attached to the second joint of the index finger (Vicon Inc.). Before neural recording, animals were trained to achieve a success rate of over 90% for performing the manual interception, to ensure a high efficiency in the arm movement and to help the animal comprehend the brain-controlled interception.

For both monkeys, two 96-channel microelectrode arrays (1.5 mm-long electrode shanks, Blackrock Microsystems) were implanted into primary motor cortex (M1) and dorsal premotor cortex (PMd) in the hemisphere contralateral to the utilized hand. The neurosurgery for implantation was operated under anesthesia (induced by 10 mg/kg ketamine, then sustained by 2% Isoflurane). Neural activities were recorded via a Cerebus acquisition system (256-channel recording system, Blackrock Microsystems), sampled at 30 kHz.

All procedures were approved by the Institutional Animal Care and Use Committee (IACUC) of the Chinese Institute for Brain Research, Beijing.

### Interception paradigm and feedforward BMI design

The arm and brain-controlled conditions were organized in blocks. The manual interception task has been recently reported (Zhang et al., 2025). At the beginning, a green disk with a radius of 1.5 cm appeared in the center of the touchscreen. When the monkey held within this central disk for 0.3 s, another 2.5 cm green disk (target) appeared at a random location, moving along an invisible circular path with a radius of 12 cm around the center disk. Conditions in which the target moved clockwise (CW: −120° /s, −240° /s) or counterclockwise (CCW: 120° /s, 240° /s), or stayed stationary (0° /s) were pseudo-randomly interleaved trial by trial. The probabilities of each target speed were equal. The central disk turned dark as a GO cue after a random delay (0.4-0.6 s); then the monkey was required to initialize an interception within 0.5 s. Once any peripheral location was touched, the target stopped. If the touched peripheral location was within a 2-cm radius of the moving target and held for 0.3 s, a red disk with the same radius as the target would appear in the touched position as feedback of success, and a drop of liquid food would be delivered. If the touch was out of the tolerance window, a blue disk appeared at the touched location as feedback of failure.

The brain-controlled interception was designed to resemble the manual interception except that monkey’s body was restricted in the monkey chair. At first, three objects appeared in the touchscreen simultaneously: a gray cursor in the screen center with a radius of 0.5 cm, a white ring with a radius of 3 cm located around the central disk, and a 1.5 cm green disk (target) on an invisible circle (10 cm radius around the center) either revolving in a constant circular motion at ±120 ° /s) or remaining static. The probabilities that the target moved at angular velocity of 120° /s, −120° /s, or 0° /s were 0.4, 0.4 and 0.2, respectively. After one second, the center cursor’s color changed from gray to white, indicating that the cursor could be driven by neural activity. The monkey was required to control the cursor to reach the ring within the premovement epoch of 0.3 second which approximated the reaction time of monkeys’ manual interception. Once the cursor crossed the ring, its color turned red, indicating the phase of ballistic flight without online correction, and it launched in the straight-line from the ring center to the crossed location, following a bell-shaped speed profile defined in equation (1) (Kao et al., 2021):

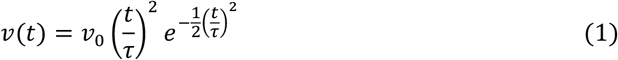

where cursor speed *v* at time *t* was adjusted based on its speed when crossing the ring *v*_0_, and *τ* was a time constant which determined the flight duration from the ring to the peripheral circle. At each time step, the distance between the cursor and target was computed. Once within 5 cm, both cursor and target halted for 0.6 s, triggering a juice reward. If the cursor was launched to the border of tolerance, the cursor would stop to move and maintain static for 0.6 s. If the cursor reached the target’s invisible circular track before a successful interception, it turned white, granting the monkey a 1 s compensatory pursuit period. Success during this period (defined by re-entering the 5 cm proximity) caused the target to turn red, a 0.6 s halt, and reward delivery. Otherwise, the trial ended without reward. Note that the cursor was not forced to stop moving at the same distance, leading to different length of launching trajectory.

In both conditions, the continuous movement direction of arm or cursor was discretized by assigning it to the central angle of its corresponding 45° sector, resulting 8 movement directions (22.5°, 67.5°, 112.5°, 157.5°, 202.5°, 247.5°, 292.5°, and 337.5°).

### Feedforward BMI training

A training protocol that included three steps was designed for the brain-controlled interception. In the first step, the monkeys were trained to perform brain-controlled center-out tasks with a delay epoch, but without ring or ballistic flight. The GO cue in the first step is the switch of cursor color from grey to white. Training began from a one-dimensional center-out task, in which a green 1.5-cm target appeared only in the left or right 10 cm from the center of the screen, and the cursor could be horizontally moved by neural activity. Once the distance between the cursor and target was less than 5 cm, the cursor would turn red as a cue of success. It was considered as a failure if the cursor reached the edge of the screen or did not reach the target in 10 seconds. Both monkeys spent about 1-2 days to achieve an accuracy rate of over 90%, then the task was switched to a two-dimensional center-out task in which the target appeared in a random location on an invisible circle with a radius of 10 cm, with the same requirement. It took both monkeys about 3-5 days to achieve an accuracy rate of over 90% in the two-dimensional center-out task.

In the second step, the monkeys were trained to perform the center-out task via the feedforward BMI (feedforward center-out). This task was one certain condition of the brain-controlled interception where the target was static. It took both monkeys about 1-2 days to achieve a success rate of over 90%.

In the final step, every session began with the two-dimensional center-out task without ring. If the success rate was over 80% after the first minute, the task switched to the feedforward center-out. If the monkey succeeded over 80% after one minute, the task switched to the feedforward interception paradigm in which the angular velocity of target was the same as the maximum in the previous day. Every time when the success rate was over 70% in 30 minutes, 5° /s would be added to the angular velocity until its absolute value reached 120 ° /s. The angular velocities of target started from 15° /s, −15° /s and 0° /s randomly interleaved in the first day of step 3. It took both monkeys about two weeks to finally intercept targets with angular velocities of ±120° /s with a success rate approaching 70%.

### Decoder initialization and calibration

The brain-controlled task was performed under the restriction of the monkey’s hands. The BMI framework for online collection of neural data, decoding and presenting decoded results was based on our ContralIt platform (Yang and Li et al. 2025).

Two asynchronous processes were used to retrieve the cursor’s motion state from the operating system and neural data from the acquisition system. The cursor’s motion state at time *t* was organized into a vector as follows:

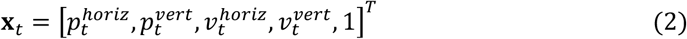

where 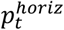 and 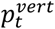 were the horizontal and vertical components of the cursor’s position, 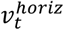 and 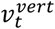 were the horizontal and vertical components of the cursor’s velocity, and constant 1 for allowing a fixed offset. The neural data at time *t* was a vector as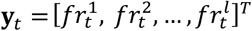, where the *l* represents the *l*th channel neural firing rate (*fr*). The BMI system reads the vector of cursor’s motion state **x**_t_ and the neural vector **y**_*t*_ every 0.02 s, paired them into a piece of data, and sent it into a first-in, first-out (FIFO) buffer with a capacity of 500 entries. The decoder was updated when this buffer was full. In the present study, recalibrated feedback intention-trained Kalman filter (ReFIT-KF) (Gilja et al., 2012) was used with the two equations:

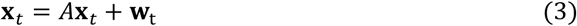

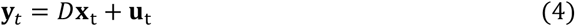

where *A* ∈ ℝ^*l*×*l*^ was the cursor’s state transition matrix, *D* ∈ ℝ^*m*×*l*^ was the observation matrix, *l* was the vector length of **x**_*t*_, *m* was the vector length of **y**_*t*_, **w**_*t*_ and **u**_*t*_ were additive Gaussian noise with zero mean and covariances *W* (state) and *U* (observation), respectively. The decoder’s parameters were updated in accordance with the following equations:

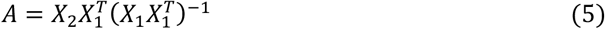

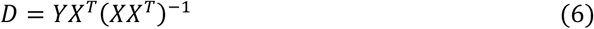

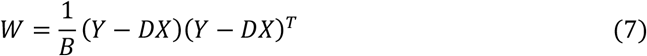

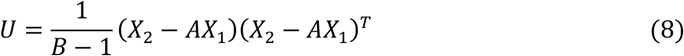

Every piece of data in the buffer was correspondingly transformed into matrix *X* ∈ ℝ^*N*×*m*^and *Y* ∈ ℝ^*N*×*l*^. For a time-series of length *N* + 1, we defined *X*_1_ covering time steps from 0 to *N* − 1 and *X*_2_ from 1 to *T. B* was the number of paired **x**_*t*_ and **y**_*t*_ in the buffer. The decoder’s parameters were continuously updated in an asynchronous process.

During the initialization of the decoder, the monkey observed a white cursor move from the screen’s center to a target position 10 cm away with the velocity of 6 cm/s. This stage ended when the decoder was updated for the first time, yet yielding the suboptimal performance.

During the decoder’s recalibration, the cursor’s velocity vector in ***x***_*t*_ was replaced by the monkey’s movement intention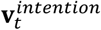, which came from equation (9).

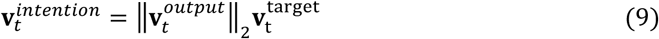

where∥·∥ denoted the *L*_2_ norm, 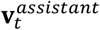 was the unit vector pointing to the target from the screen center and 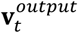 followed equation (10).

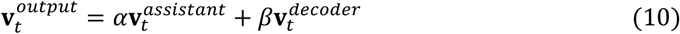

where 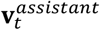 was the assistance vector which was a unit vector and always pointed to the target from the screen center, 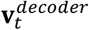 was the decoder vector which was the output of trained Kalman filter, and *α* and *β* were adjustable weights. 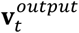 was to guide the monkey’s performance above a threshold by adjusting parameters during the recalibration. It was assumed that a movement intention pointing to the target was continuously generated by the monkey. In this stage, the BMI system kept collecting movement intention and updating the decoder’s parameters. Specifically, the BMI system operated as follows: when the monkey performed the one or two-dimensional center-out task, *α* would decrease by 1 from 6 while *β* would increase by 0.2 from 0 each time when the success rate exceeded 90% after one minute. When *α* decreased to 1, the decrease stride of *α* was changed to 0.1. Eventually, when *α* decreased to 0, and *β* increased to 1.5, and the success rate exceeded 90% after one minute, the decoder’s update stopped, and we started to collect data with fixed parameters over 30 minutes. Then the monkey began to perform the feedforward center-out task, the ring appeared and *β* decreased to 1. When the success rate was higher than 70% after one minute, the task paradigm switched to interception. When the success rate was higher than 70% after one minute, the recalibration was finished, and we started to collect data.

### Feedforward BMI performance evaluation

Three performance metrices for BMI were used in the present study: success rate, path efficiency, and time to target. The success rate was the percentage of successful-interception trials. The path efficiency was calculated according to equation (11):

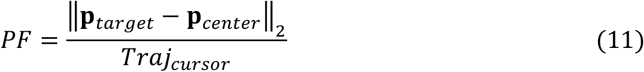

where ∥·∥ denoted the *L*_2_ norm, **p**_*target*_ was the position vector of the target, **p**_*center*_ was the position vector of the screen center, and the *Traj*_*cursor*_ was the length of the cursor’s trajectories in a trial. Time to target was defined as the time from the start of brain control to when the cursor reached the target. All the performance metrices were calculated in one session with the exclusion of trials during the decoder’s initialization and recalibration.

### Offline processing

In the present study, 34 sessions (20 brain-controlled conditions and 14 arm conditions) were recorded in monkey B, and 44 (25 and 19) for monkey L. Spikes were sorted via Kilosort 2.5 (Pachitariu et al., 2024). For monkey B, 74±7 units were well-isolated for brain-controlled and 94±29 for manually-controlled conditions. For monkey L, the corresponding counts were 145±14 and 174±34.

The Kilosort 2.5 algorithm was deployed in the singularity container provided by the spikeinterface (Buccino et al., 2020). All the data was offline sorted and checked manually. The sorted neural data and behavioral data were organized according to the Neurodata Without Borders (NWB) data standard (Rübel et al., 2022) and then loaded and analyzed with Pynapple (Viejo et al., 2023).

### Neural representation analysis

The peri-event time histograms (PETHs) of each unit were calculated with spike trains aligned to corresponding event, binned into 0.02 s window and then smoothed with a Gaussian kernel (SD=0.06 s). Firing rates were averaged across trials according to the eight movement-directions (22.5°, 67.5°, 112.5°, 157.5°, 202.5°, 247.5°, 292.5°, and 337.5°), and the preferred direction in each bin (0.02 s) was estimated using:

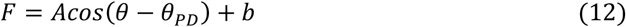

where *F* was the firing rate, *A* the tuning depth, *θ* the arm movement direction or the cursor’s ballistic direction, *θ*_*PD*_ the estimated preferred direction, and *b* the baseline. |Δ*θ*_*PD*_|, the absolute difference of preferred direction between two epochs, was defined as follows:

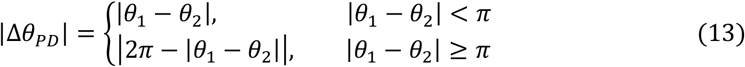

where *θ*_1_and *θ*_2_ were preferred directions in two epochs.

### Neural activity decomposition analysis

We constructed a dataset of neural population activities. All neurons’ firing rates were preprocessed as described earlier in neural representation analysis, normalized as proposed by Churchland et al. (Churchland et al., 2012), demeaned by subtracting the trial-averaged mean, then averaged according to the arm movement or ballistic directions to obtain a matrix of *N* ∈ ℝ^*k*×*n*×*t*^, where *k* denoted the number of movement direction, *n* the number of neurons, and *t* the number of time bins. Then we reshaped *N* into matrix *M* ∈ ℝ^*k*×*nt*^, performed principal component analysis (PCA) on *M* by extracting the PC from *nt* dimension and selected the top three PCs to construct the feature subspace. Neural trajectories were obtained by projecting the vectors of matrix *N* in the same arm movement or ballistic direction in each time point onto the space and then concatenating them.

The overlapping extent of premovement and execution subspaces was estimated by the alignment index (Elsayed et al., 2016). We took two sections from matrix *M* to cover the premovement epoch (*M*_*prem*_, −0.3 s to −0.28 s to MO) and the execution epoch (*M*_*exec*_, 0.18 s to 0.2 s from MO). We performed PCA on *M*_*prem*_ and *M*_*exec*_, and selected their top ten PCs to construct the premovement subspace *C*_*prem*_ and execution subspace *C*_*exec*_, respectively. The alignment index between these two subspaces was calculated according to:

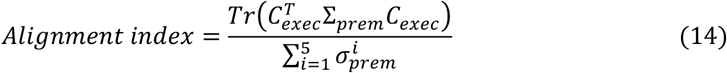

where Σ_*prem*_ denoted the covariance matrix of premovement neural activity *M*_*prem*_, and 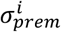 denoted the *i*th singular value of *C*_*prem*_. Population activities in the preparatory and execution epochs were projected onto the Stiefel manifold with 6 dimensions, and the corresponding subspaces of this manifold were computed using pymanopt. Variances of population activities captured during the preparatory and execution epochs were calculated by projecting these activities onto the corresponding subspaces. The alignment indexes of the control group in Fig. 3d and h were from 20 sets of data, randomly generated according to Elsayed et al. proposed (Elsayed et al., 2016).

### jPCA projection

In the present study, jPCA was used to identify low-dimensional rotational structure in the neural population dynamics (Churchland et al., 2012). Spike trains aligned to movement onset (manual condition: −0.05 s to 0.15 s, brain-control condition: −0.1 s to 0.3 s) were processed as follows: binned into 0.02 s windows, smoothed with a Gaussian kernel (SD = 0.06 s), averaged by discretized movement direction, normalized and reshaped into matrix *S* ∈ ℝ^*k*×*nt*^ (*k* the number of movement direction, *n* the number of neurons, *t* the number of time bins). In jPCA analysis (https://churchland.zuckermaninstitute.columbia.edu/content/code), we fitted a matrix *M*_*skew*_ using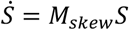, yielded two eigenvectors with the largest imaginary eigenvalues (*V*_1_ and *V*_2_) and then two orthogonal vectors jPC_1_ = *V*_1_ + *V*_2_ and jPC_2_ = *j*(*V*_1_ + *V*_2_), and projected neural activity *S* onto the two-dimensional subspace spanned by jPC_1_ and jPC_2_. In BMI condition, we fitted neural firing rate to cursor velocity direction in each 0.02 s time bin. If the *R*^2^ of a neuron was larger than 0.8 in over half of time bins, this neuron was believed to represent the movement state of cursor. These types of neurons were selected and concatenated across all datasets collected from BMI condition for extracting jPC and plotting the neural trajectory in jPCA space. In the arm condition, we selected two example datasets, called ‘20240920’ and ‘20250120’ from monkey B and L, respectively.

### Canonical correlation analysis

The firing rates in manual and brain-controlled conditions were calculated from spike trains which were aligned to movement onset (−0.1 s to 0.3 s), binned into 0.02 s windows, and smoothed with a Gaussian kernel (SD = 0.06 s). In the canonical correlation analysis (CCA), the firing rates were organized into matrixes *N*_*hand*_ and *N*_*cursor*_, yielding *Q*_*hand*_ and *R*_*hand*_, *Q*_*cursor*_ and *R*_*cursor*_ by the QR decomposition. The across-condition covariance 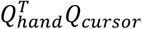 yielded *U*ΣV^*T*^ via singular value decomposition (SVD), this further yielded the aligned neural manifolds 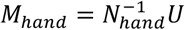 and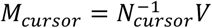. The aligned neural data, resulting from projecting matrixes *N*_*hand*_ and *N*_*cursor*_ onto their corresponding aligned neural manifolds, were used for finding the canonical correlation in the different neural modes (Safaie et al., 2023).

## Supplementary

**Supplementary Fig. S1.**
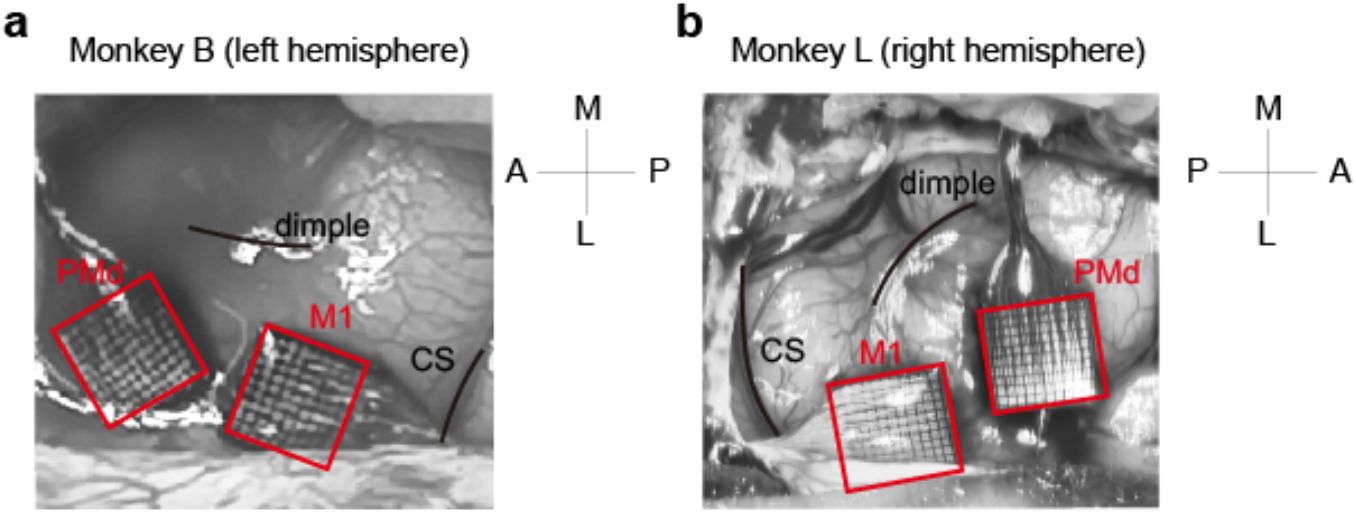
Implantations of Utah arrays into the motor cortex. Two Utah arrays implanted into the dorsal premotor cortex (PMd) and primary cortex (M1), respectively, were marked with red squares in the pictures. Central sulcus and dimple were marked with black lines to indicate the referenced landmarks for implantation.

**Supplementary Fig. S2.**
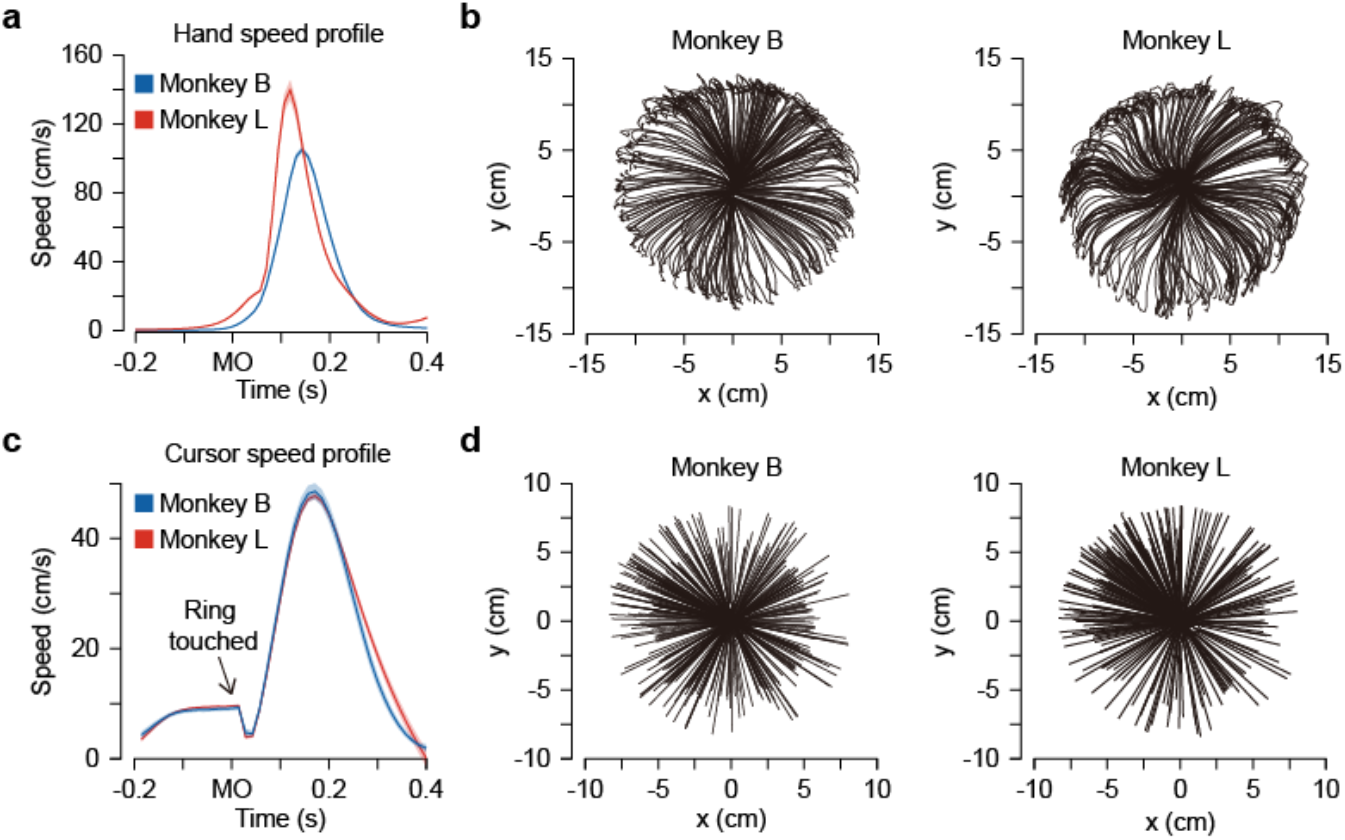
Trajectories of hand and cursor. (**a**) A marker on one finger of the monkey was labeled to track the trajectories of hand movement. Time point ‘0’ was aligned to the onset of arm movement. (**b**) Hand movement trajectories in the manual interception task. Time window: from the Go-cue onset to target touching. (**c**) Ballistic velocity profile of the cursor. Time point ‘0’ was aligned to the time when the cursor touched the ring. (**d**) Cursor movement trajectories in feedforward BMI task. Time window: from the onset that the cursor could be operated to the time when the cursor touched the target.

**Supplementary Fig. S3.**
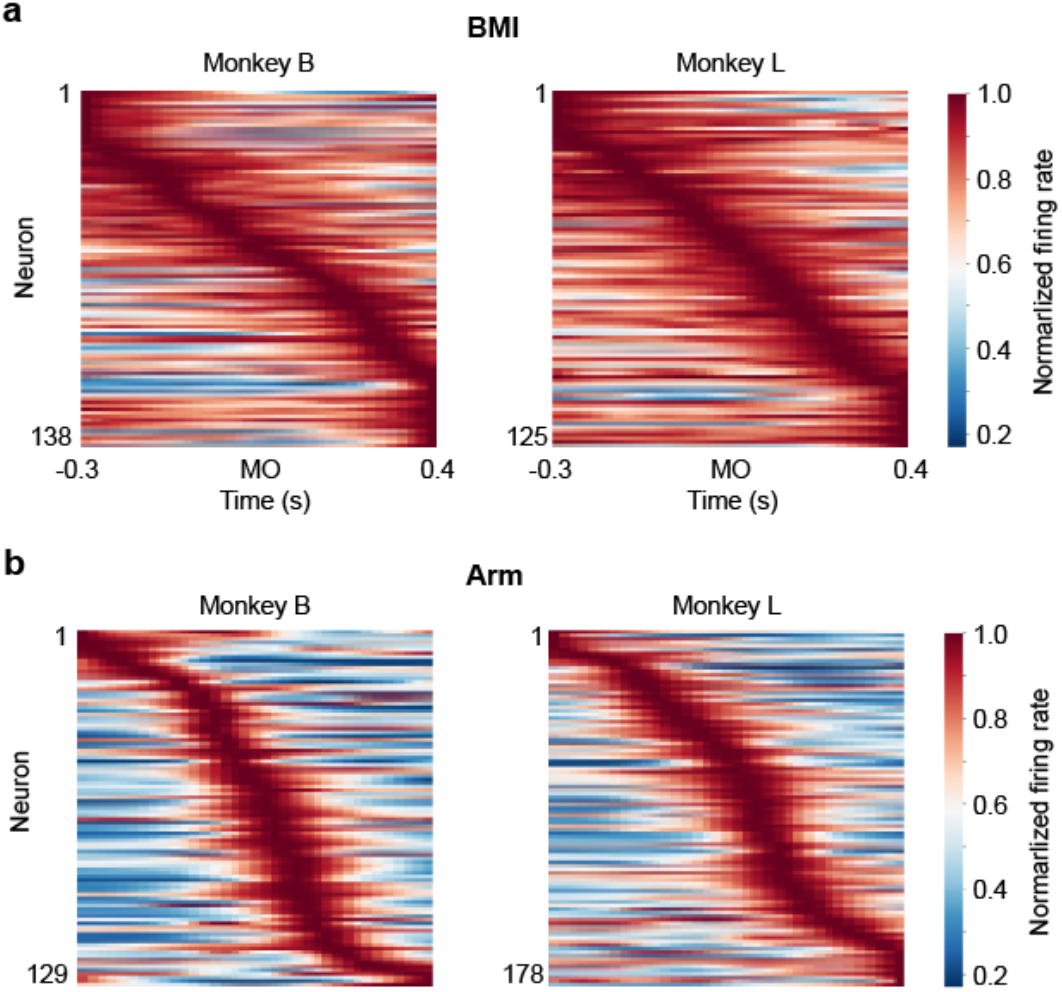
Heatmaps of normalized firing rates after being averaged across all the trials for single units in the BMI condition (**a**) and arm condition (**b**) for both monkeys. Neural activities were aligned to movement onset (MO), with units sorted with the time of peak firing rate.

**Supplementary Fig. S4.**
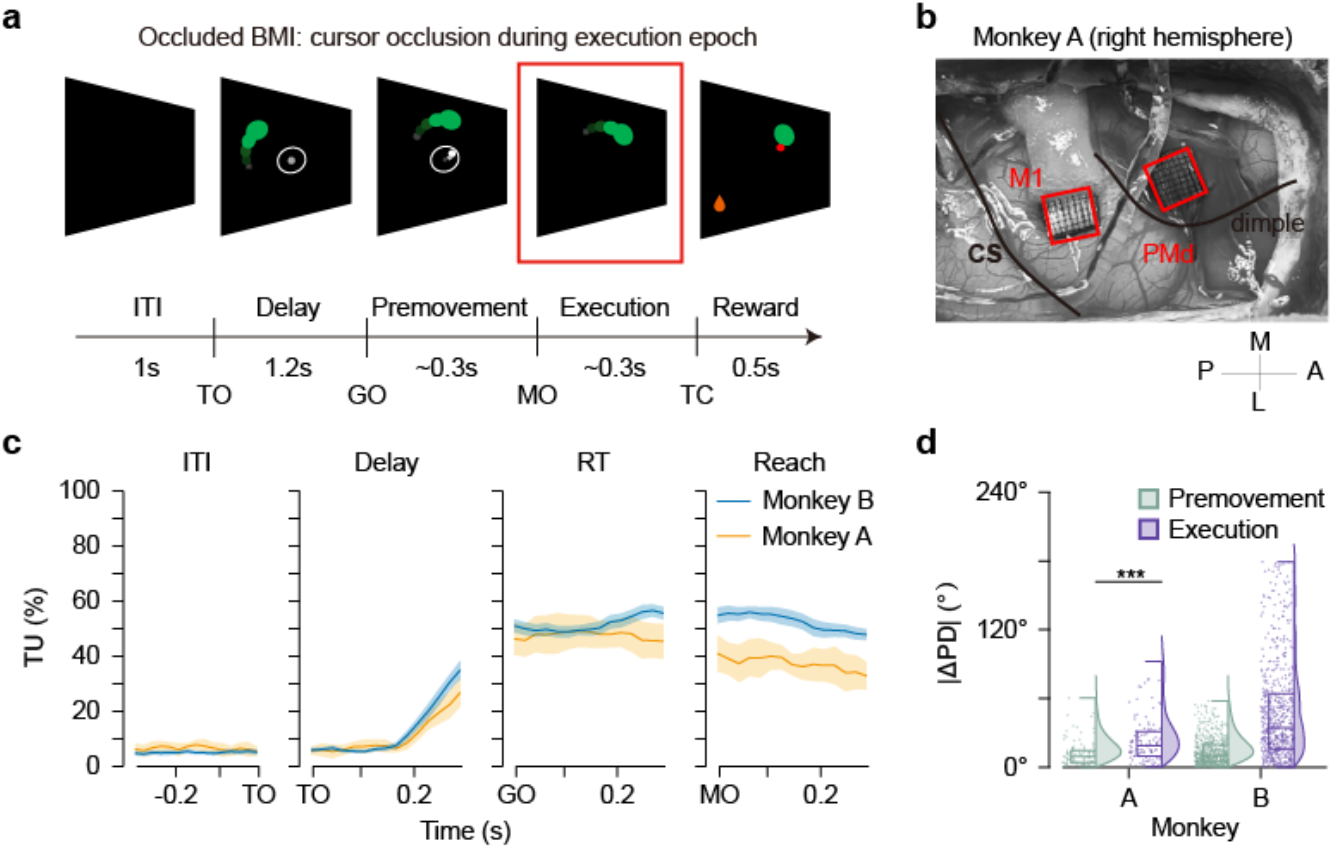
Tunings of single units under the occluded BMI condition (i.e. ballistic cursor was occluded in the execution epoch). (**a**) The schematic of the occluded BMI paradigm, where cursor disappeared during its ballistic flight epoch (highlighted by the red box). (**b**) Implantations of Utah arrays into the motor cortex of a new monkey, A. (**c**) Tuned unit’s percentage (TU%) in different epochs when aligned to the onset of target on (TO), Go-cue on (GO), and movement onset (MO) in the occlusion condition. (**d**) Absolute differences of PD between premovement and delay epochs (green) and between execution and delay epochs (purple) for both monkeys.

**Supplementary Fig. S5.**
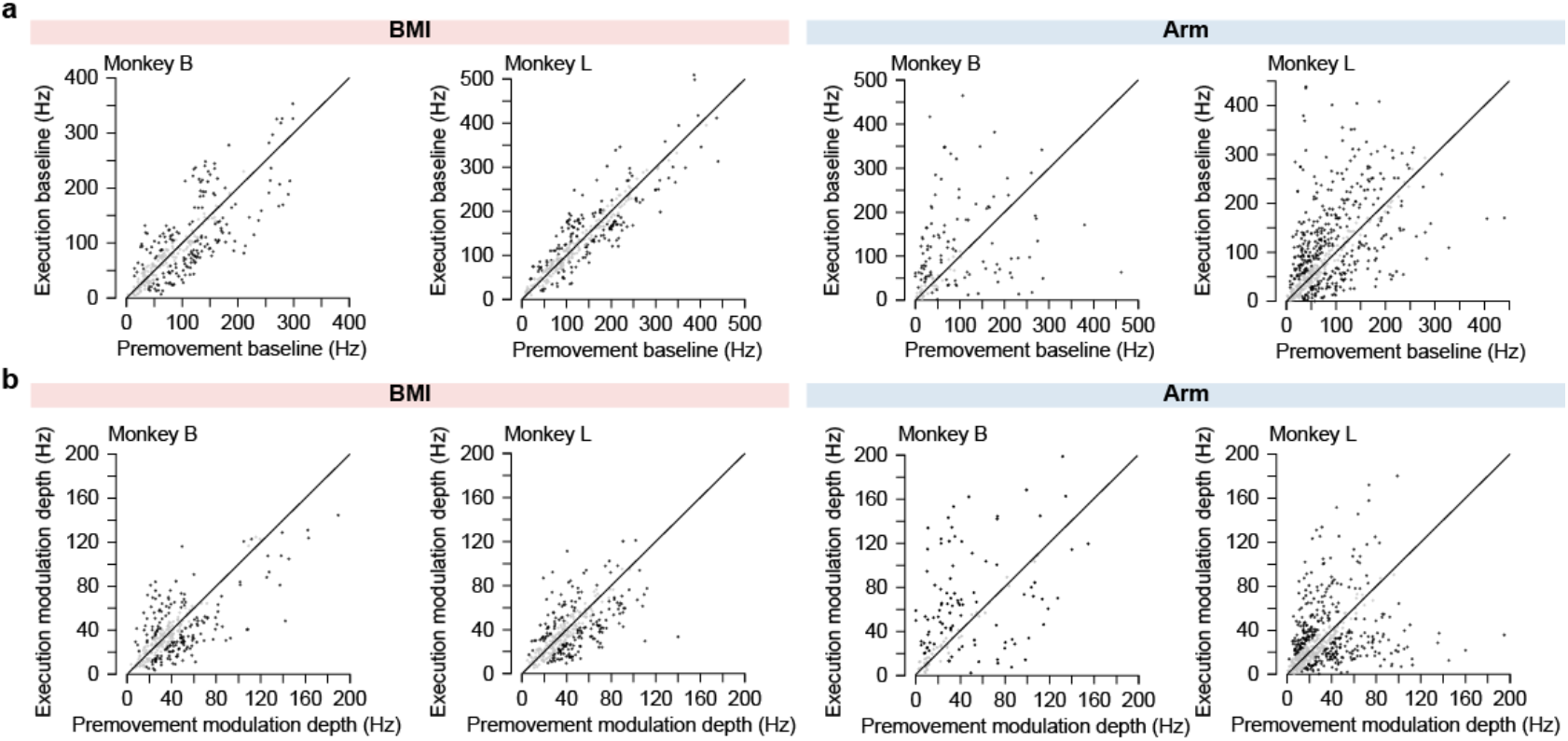
Firing rates of ‘baseline’ neurons (**a**) and ‘modulation depths’ neurons (**b**) in premovement epoch (−200 ms to −180 ms, aligned to MO) and execution epoch (180 ms to 200 ms, aligned to MO) in the BMI and arm conditions for both monkeys. Dark dots represent neurons with significant change of baseline or modulation depth.

**Supplementary Fig. S6.**
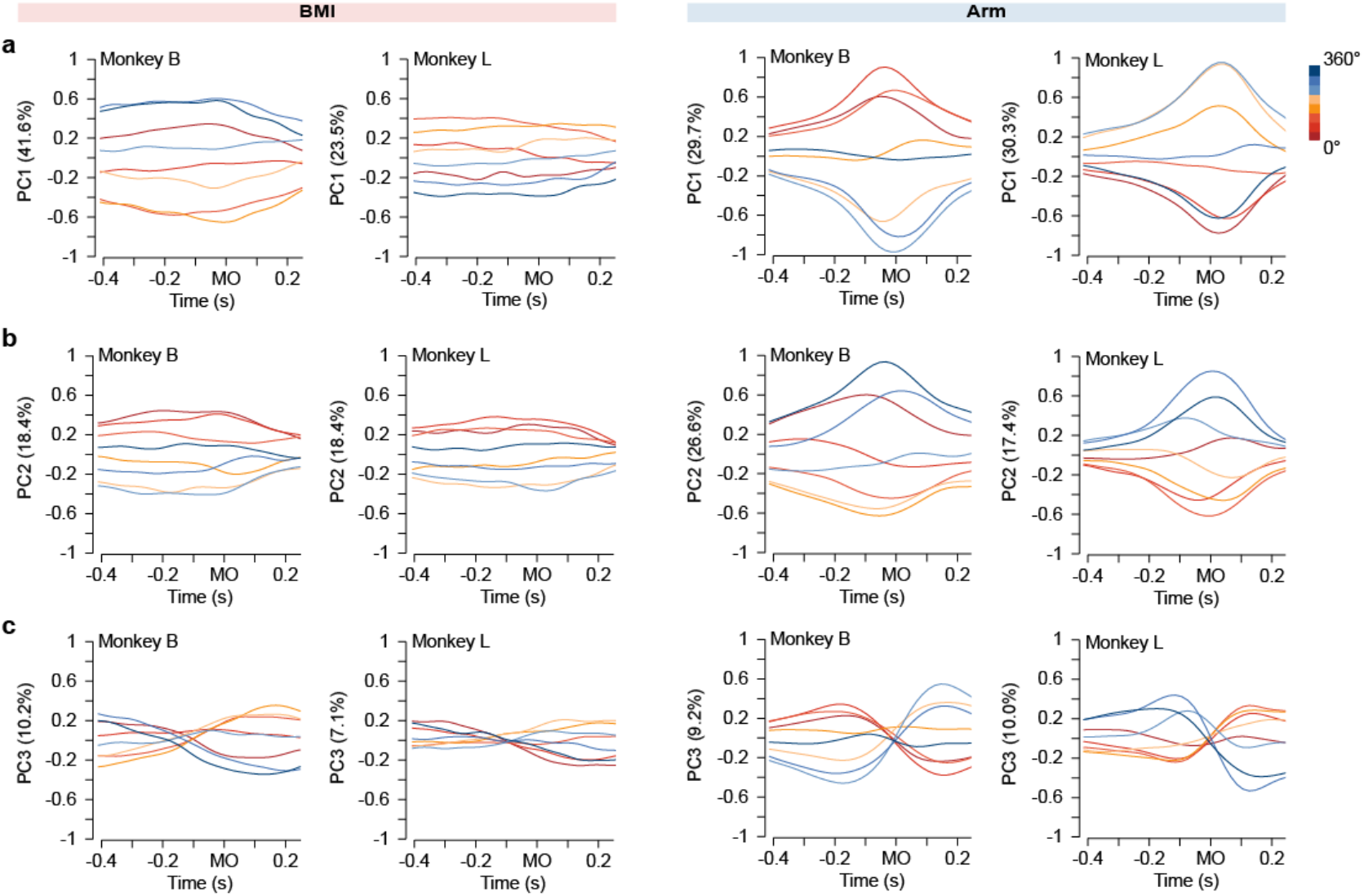
Top three PCs in eight movement directions. Colors represent different movement directions.

